# A Phylogenomic Assessment of Processes Underpinning Convergent Evolution in Open-Habitat Chats

**DOI:** 10.1101/2022.06.21.496980

**Authors:** Niloofar Alaei Kakhki, Manuel Schweizer, Dave Lutgen, Rauri C. K. Bowie, Hadoram Shirihai, Alexander Suh, Holger Schielzeth, Reto Burri

## Abstract

Insights into the processes underpinning convergent evolution advance our understanding of the contributions of ancestral, introgressed, and novel genetic variation to phenotypic evolution. Phylogenomic analyses characterizing genome-wide gene tree heterogeneity can provide first clues about the extent of ILS and of introgression and thereby into the potential of these processes or (in their absence) the need to invoke novel mutations to underpin convergent evolution. Here, we were interested in understanding the processes involved in convergent evolution in open-habitat chats (wheatears of the genus *Oenanthe* and their relatives). To this end, based on whole-genome resequencing data from 50 taxa of 44 species, we established the species tree, characterized gene tree heterogeneity, and investigated the footprints of ILS and introgression within the latter. The species tree corroborates the pattern of abundant convergent evolution, especially in wheatears. The high levels of gene tree heterogeneity in wheatears are explained by ILS alone only for 30% of internal branches. For multiple branches with high gene tree heterogeneity, D-statistics and phylogenetic networks identified footprints of introgression. Finally, long branches without extensive ILS between clades sporting similar phenotypes provide suggestive evidence for a role of novel mutations in the evolution of these phenotypes. Together, our results suggest that convergent evolution in open-habitat chats involved diverse processes and highlight that phenotypic diversification is often complex and best depicted as a network of interacting lineages.

## Introduction

Molecular phylogenetics has unveiled many previously unknown examples of convergent evolution – here meant to refer to a *phenotypic pattern* in which non-sister species are phenotypically more similar to each other than to their respective sister species (following Arendt and Reznick 2008; Stern 2013). Under such an evolutionary outcome, species relationships based on morphometrics, coloration, behavior, or other ecological traits are discordant with the history of descent reflected in the species tree (Aliabadian et al. 2012; Elmer and Meyer 2011; Jarvis et al. 2014; Martin and Orgogozo 2013; Paterson et al. 2020; Schweizer et al. 2019a; Schweizer et al. 2019b; Stern 2013). While the many observations of such discordances across the tree of life witness of the abundance of convergent evolution, insights into the underlying processes remain more elusive.

Phylogenetic information from genomic data now provides unprecedented power to consolidate patterns of convergent evolution and obtain insights into the underlying processes. Many examples of putative convergent evolution are yet based on phylogenies reconstructed from a restricted number of genetic markers (Aliabadian et al. 2012; Brusatte et al. 2015; Colosimo et al. 2005; Cresko et al. 2004; Stern 2013). Since phylogenetic relationships at different positions in the genome, referred to as ‘gene trees’, can vary substantially, many gene trees inevitably deviate from the species’ history of descent reflected in the species tree (Degnan and Rosenberg 2006; Toews and Brelsford 2012). Hence, the mismatch of single gene trees with phenotypic similarities alone does not provide conclusive evidence for convergent evolution (Degnan and Rosenberg 2006; Doyle 1997; Lamers et al. 2012). Confirming instances of convergent evolution, therefore, requires species tree reconstructions from genome-wide variation. Once the evidence for convergent evolution is corroborated by the species tree, we can move forward to investigate the processes underlying gene tree heterogeneity that may also underpin convergent evolution.

Convergent evolution can occur via three processes: First, phenotypic similarities can evolve through independent mutations in the same or different genes (“parallel evolution” *sensu* Stern 2013) (Arendt and Reznick 2008; Martin and Orgogozo 2013; Stern 2013). In Mexican cavefish (*Astyanax mexicanus*), for instance, the evolution of decolored brown phenotypes and albinism in separate caves occurred through different mutations in the MC1R and OCA2 genes (Gross et al. 2009; Protas et al. 2006; Stahl and Gross 2015). Similarly, in plants, isoforms of PEPC found in C4 photosynthesis, and similar floral traits important for pollination have evolved multiple times independently (Besnard et al. 2009; Christin et al. 2007; Hoballah et al. 2007; Preston and Hileman 2009; Whittall et al. 2006).

The second, and likely most frequent process leading to convergent evolution that also accounts for most gene tree heterogeneity is incomplete lineage sorting (ILS; Stern 2013 includes this under “collateral evolution”), that is, the retention of alleles and traits that were already present in the ancestral lineage (Colosimo et al. 2005; Cresko et al. 2004; Stern 2013; Van Belleghem et al. 2018). ILS is prevalent in radiations characterized by large effective population sizes and fast succession of speciation events, such as in the evolution of neoavian birds (Jarvis et al. 2014; Suh 2016; Suh et al. 2015) or in the diversification of sticklebacks (Colosimo et al. 2005; Jones et al. 2012; Roberts Kingman et al. 2021). In such cases, a high proportion of ancestral variation may be retained over subsequent species splits and segregate in the independently segregating gene pools of daughter species (Maddison 1997). Selection or drift in non-sister species may fix the same genotype (and phenotype), while sister species may fix a different genotype/phenotype. For instance in Humans 1% of the genome is genetically more similar to orangutans than to chimps due to ILS, even though these primates are characterized by small effective population sizes (Hobolth et al. 2011).

Third, in hybridizing lineages, convergent evolution and gene tree heterogeneity may be underpinned by introgression (the exchange of genetic material between species) that mingles genotypes and phenotypes among species (Stern 2013 includes this under “collateral evolution”) (Heliconius Genome Consortium 2012; Malinsky et al. 2018; Song et al. 2011; Stryjewski and Sorenson 2017). In particular, introgression between non-sister species may result in these species being phenotypically more similar than they are to their respective sister species, such as exemplified by wing-pattern mimicry in *Heliconius* butterflies (Edelman et al. 2019; Pardo-Diaz et al. 2012), and by plumage coloration of *Munia* finches and of members of the Black-eared Wheatear (*Oenanthe hispanica*) complex (Schweizer et al. 2019a; Stryjewski and Sorenson 2017). Importantly, in an increasing number of instances, such as in *Heliconius* butterflies, Yellowstone wolves, Darwin’s finches, and cichlid fish, introgression has exchanged alleles between species and resulted in the formation of beneficial phenotypes (Enciso-Romero et al. 2017; Genner and Turner 2012; Grant et al. 2005; Lamers et al. 2012; Wallbank et al. 2016). Given that over the last decade genomic studies have contributed increasing evidence for the abundance of such adaptive introgression, hybridization (the interbreeding of different species) may underpin convergent evolution more often than previously appreciated (Campagna et al. 2017; Han et al. 2017; Marques et al. 2019a; Meier et al. 2018).

Multiple factors influence which of these processes were most likely involved in specific cases of convergent evolution. These factors include the evolutionary time scale under consideration, the speed at which successive speciation events occurred, effective population sizes, and the opportunity for genetic exchange according to biogeographic history. Waiting times for beneficial mutations are long (Barrett and Schluter 2008; Hedrick 2013; Hermisson and Pennings 2005). Independent mutations with the same phenotypic effect are thus exceedingly rare (Eyre-Walker and Keightley 2007) and only over the course of millions of years may occur in sufficient number to be a source of convergent evolution (Hedrick 2013). Therefore, at short evolutionary time scales, convergent evolution may more often involve the recruitment of standing genetic variation (Barrett and Schluter 2008), notably from the pool of ancestral variation segregating in extant species, or variation introgressed from other species (Stern 2013); especially since in young lineages ancestral variation is still segregating and because reproductive isolation may still be incomplete between young species.

Phylogenomics can provide important indirect insights into the potential contribution of ILS, introgression, and novel mutations to convergent evolution: First, the species tree provides initial clues on whether speciation events occurred over short enough time scales for ancestral variation to be passed to descent lineages and thus remain incompletely sorted in important proportions beyond speciation events. Second, insights into the extent of ILS and presence of introgression can be gained from levels of gene tree heterogeneity (Degnan and Rosenberg 2006; Funk and Omland 2003; Jarvis et al. 2014; Nater et al. 2015; Suh 2016; Suh et al. 2015) and symmetries of gene tree frequencies (Hibbins and Hahn 2022). Gene tree heterogeneity is high under both ILS and introgression, but the two processes leave different proportions of alternative gene trees, based on which they can be distinguished (Hibbins and Hahn 2022; Sayyari and Mirarab 2018; Sayyari et al. 2018). In the presence of extensive ILS or of introgression, a parsimonious approach attributes the source of convergent evolution to these processes, even though independent mutations cannot be excluded as the source of convergent evolution (Colosimo et al. 2005; Cresko et al. 2004; Pardo-Diaz et al. 2012; Stryjewski and Sorenson 2017). The absence of detecting these processes, conversely, would indirectly suggest novel mutations as a potential source of convergent evolution. Therefore, surveys of gene tree heterogeneity and symmetries of gene tree proportions represent a promising avenue to probe the potential of the alternative processes to contribute to convergent evolution.

Here, we reconstructed the species tree and assessed the contribution of ILS and introgression to gene tree heterogeneity in open-habitat chats (genera *Campicoloides, Emarginata, Myrmecocichla, Oenanthe, Pinarochroa* and *Thamnolaea*), a monophyletic group of songbirds displaying a high incidence of convergent evolution (Aliabadian et al. 2012; Mayr and Stresemann 1950; Schweizer et al. 2019a; Schweizer et al. 2019b). The phylogenetic relationships among open-habitat chats inferred from mitochondrial data were entirely unexpected from a morphological perspective (Aliabadian et al. 2012). Species similar in plumage coloration and other traits were often spread far apart across the mitochondrial phylogeny, suggesting convergent evolution of phenotypic similarities (Aliabadian et al. 2012; Outlaw et al. 2010; Schweizer and Shirihai 2013; Schweizer et al. 2019a; Schweizer et al. 2019b). For a limited subset of species studied, genome-wide variation (ddRAD data) confirmed the mitochondrial relationships (Schweizer et al. 2019b). Furthermore, hybridization resulted in substantial introgression in the *Oenanthe hispanica* complex (Schweizer et al. 2019a) and is suspected to have played a role in phenotypic and species evolution in the *O. picata* complex (Panov 2005). In these instances, introgression between non-sister taxa may well explain convergent evolution. However, genomic data is essential to corroborate and refine the species tree and assess the incidence of ILS and/or introgression across open-habitat chats.

Based on whole-genome resequencing data from 50 taxa of 44 open-habitat species (**Tab. S1**), we aimed to obtain insights into the potential roles of alternative processes in driving convergent evolution in these songbirds. To this end, we (i) reconstructed the species tree, (ii) estimated gene tree variation across the genome, and (iii) explored ILS and introgression as drivers of the underlying high gene tree heterogeneity. Our results reveal a comprehensive picture of open-habitat chat evolution involving high rates of ILS and multiple instances of introgression particularly in wheatears (genus *Oenanthe*). Footprints of ILS and introgression as well as considerable divergence times between the main clades of wheatears with convergent evolution suggest that, most likely, a combination of ILS, introgression, and novel mutations explains the convergent evolution observed in wheatears.

## Results

### Sampling, Nuclear Data Preparation, and Mitogenome Assembly

To achieve an almost complete taxon sampling, we resequenced the genomes of 50 open-habitat chat taxa from 44 of 47 recognized species (**Fig. 1**; **Tab. S1**). A *Saxicola maurus* genome was included as outgroup (Sangster et al. 2010; Zuccon and Ericson 2010). We mapped the sequencing reads to the reference genome assembly of *Oenanthe melanoleuca* (Peona et al. in prep.) and followed GATK best practices for nuclear data preparation. Mapping efficiency was not correlated with the degree of divergence from the reference genome, but data obtained from DNA extracted off museum skins mapped at a lower percentage (linear model, d_*XY*_: t=-0.41, p=0.68; tissue_museum_: t=-6.56, p<0.001; R^2^=0.53). After mapping, sequencing coverage ranged from 4.6 x to 40.6 x, with an average coverage of 12.2 x ± 6.2 x (**Tab. S1**). We extracted mitochondrial sequence data for all 13 protein-coding genes and two rRNA genes from the resequencing data using MitoFinder 1.2 (Allio et al. 2020). To ensure that results did not depend on filtering strategy, all analyses were run with four sets of differently filtered data (see Material and Methods).

**Figure 1.**
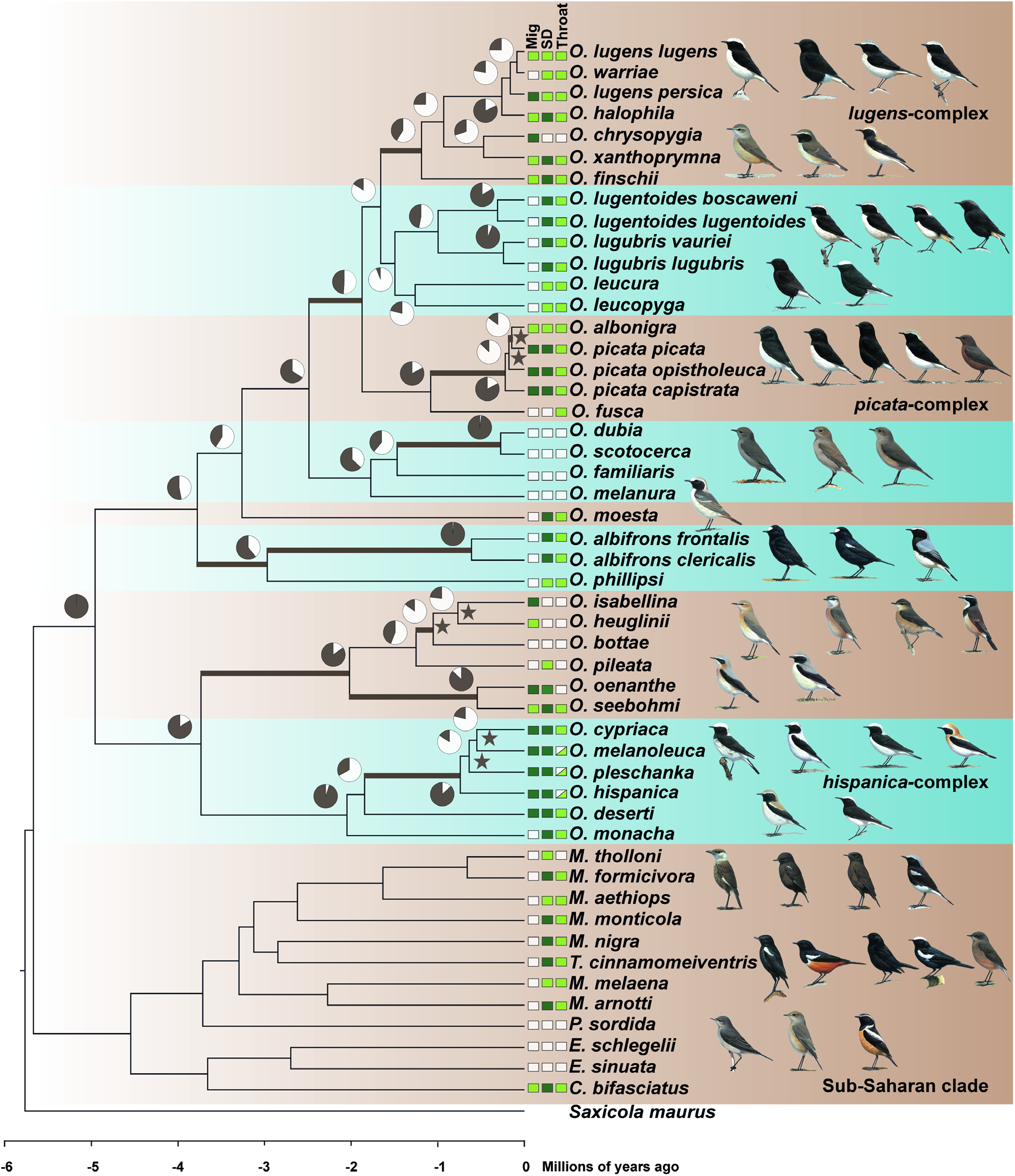
Time-calibrated phylogenetic tree of open-habitat chats and levels of ILS. All nodes are supported by bootstrap values of 100. Pie charts depict the gene tree heterogeneity for each internal branch, with the brown proportion indicating the proportion of concordant gene trees (gCF). Coloured branches indicate internal branches for which ILS alone is statistically sufficient to explain the observed gene tree heterogeneity. Stars indicate branches that are in the phylogenetic anomaly zone. The character states of three selected characters: Sexual dimorphism (SD), monomorphic female-type (white), monomorphic male-type (pale green), dimorphic (dark green); Migratory behaviour (Mig), sedentary (white), short-distance migrant (pale green), long-distance migrant (dark green); and throat coloration (throat), white (white), black (pale green), and polymorphic (white and pale green). Drawing courtesy of Chris Rose (www.chrisrose-artist.co.uk) with permission from Bloomsbury Publishing Plc.

### Species Tree Reconstruction Based on Nuclear Genomic Data

We first set out to reconstruct and root the species tree based on regions of the genome least likely affected by mapping biases. To this end, we extracted data from genomic intervals hosting avian Benchmarking Universal Single-Copy Orthologs (BUSCO). This resulted in data from 7,335 BUSCO, with alignment lengths varying from 89,898 kb to 140,640 kb (depending on filtering strategy) for ML analyses of concatenated data, respectively 2,091 BUSCO with alignment lengths varying from 10,575 kb to 15,290 kb for LD-pruned data free of interlocus recombination for multispecies coalescent-based species tree reconstruction. Results were consistent among filtering strategies. Hence, we only report results based on the most stringent filtering of read depth (ii, DP=5, PW=50%, MD=15%). Both, maximum likelihood (ML) analyses in IQtree2 based on concatenated data and multi-species coalescent analyses in ASTRAL-III (based on BUSCO ML gene trees) established sub-Saharan species of the genera *Campicoloides, Emarginata, Myrmecocichla, Pinarochroa*, and *Thamnolaea* as the sister clade to all other open-habitat chats (**Fig. S1a**). For the subsequent analyses we excluded the *Saxicola* outgroup and rooted the trees on the sub-Saharan clade.

We then moved to reconstruct the species tree based on an as broad representation of the genome as possible. To this end, we extracted alignments including variant and invariant sites for non-overlapping 10 kb windows. We henceforth refer to these windowed data as “loci”. Analyses included only loci that fulfilled filtering criteria for read depth, alignment length, data missingness (see Material and Methods), and absence of evidence for intra-locus recombination. Furthermore, we sub-sampled filtered loci to be at least 10 kb apart to ensure free inter-locus recombination. Depending on filtering strategy, this left us with 5,267-6,791 loci with total alignment lengths of 34,556-52,243 kb (**Tab. S2**). We identified branches in the “anomaly zone” (Degnan and Rosenberg 2006) in several clades of wheatears: in the *hispanica* and *picata* complexes, and in the *isabellina* clade **(Fig. 1)**. Nevertheless, the polytomy test based on local quartet supports in ASTRAL-III showed no evidence for polytomies in the species tree (P = 0 for all branches). The ML tree based on concatenated data and the multi-species coalescent-based species tree were fully supported and in agreement both with each other (except the position of T. *cinnamomeiventris* within the sub-Saharan clade) and with the tree based on BUSCO (**Fig. S1b**). Finally, a SNP-based species tree estimated in SVDquartets mostly confirmed the sequence-based results (**Fig. S2**). The only three disagreements (position of *O. leucura* and *O. leucopygia*, position of *O. bottae* and *O. pileata*, and position of *T. cinnamomeiventris*) were poorly supported in the SNP-based analysis and are likely a result of high levels of ILS under which sequence-based approaches are more accurate than approaches based on SNP data alone (Chou et al. 2015).

### Mitogenomic Relationships and Mito-Nuclear Discordances

We were interested in whether previously inferred relationships based predominantly on single mitochondrial genes (Alaei Kakhki et al. 2016; Aliabadian et al. 2012; Schweizer and Shirihai 2013) were supported by full mitogenomes and in inferring mito-nuclear discordances.

Mitogenomic relationships were in remarkable agreement with previously inferred phylogenetic relationships based predominantly on individual mitochondrial genes (Alaei Kakhki et al. 2016; Aliabadian et al. 2012; Schweizer and Shirihai 2013), yet showed several discordances with the species tree recovered from nuclear data (**Fig. 2**). Mito-nuclear discordances in wheatears were found in several places across the species tree but were mostly restricted to the placements of tip taxa: (i) In the *lugens* complex, nuclear data placed *O. I. persica* within the complex, whereas the mitogenome placed it with *O. xanthoprymna* and *O. chrysopygia.* (ii) In the *picata* complex, *O. albonigra* that by mitochondrial data was considered a sister taxon to the *picata* complex, was placed within the latter as a sister taxon to the phenotypically almost identical *O. p. picata* by nuclear data. (iii) In the *hispanica* complex, *O. cypriaca* was placed as sister to either *O. melanoleuca* or *O. pleschanka* in nuclear and mitogenomic data respectively. (iv) In the *isabellina* clade, *O. heuglini* as sister to either *O. isabelline* or *O. bottae* by nuclear or mitogenomic data, respectively. (v) Moreover, *O. leucura* and *O. leucopyga* formed a sister clade to the *O. lugubris/lugentoides* clade according to the nuclear species tree, but mitogenomes placed them consecutively at the root of the clade including *O. finschi* and the *lugens* complex. To understand whether nuclear gene trees were entirely discordant with mitogenomic relationships or in part reflected the latter, for each of the above discordances we checked for nuclear gene trees that agreed with the mitogenomic tree. This showed that for most of the mitonuclear discordances, roughly 15% of the gene trees agreed with the mitogenomic relationships (*picata* complex: 14.40%, 4,282 of 29,730 gene trees; *hispanica* complex: 13.13%, 3,905 of 29,730 gene trees; *isabellina* clade: 15.71%, 4,671 of 29,730 gene trees; *lugens* complex: 2.77%, 824 of 29,730 gene trees).

**Figure 2.**
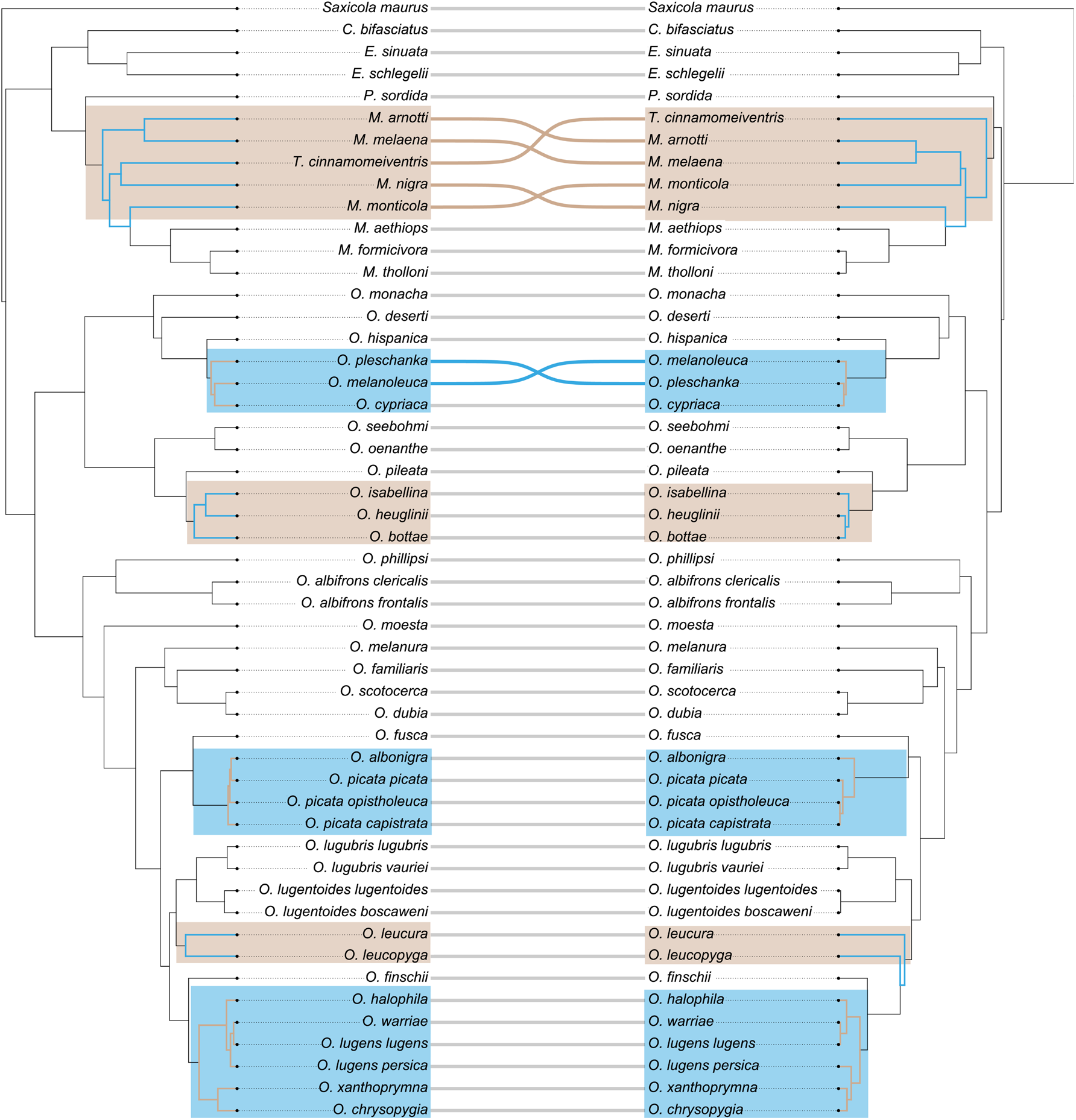
Mito-nuclear discordances. Shown are the time-calibrated phylogenetic trees based on nuclear data (left) and full mitogenomes (right).

### Time trees

In addition to species’ relationships, we were interested in understanding the time scales at which species diverged. Due to the lack of appropriate fossils, we resorted to first estimating a time-calibrated mitochondrial phylogeny based on the 13 mitochondrial protein-coding genes, for which substitution rates are available (Lerner et al. 2011). The analysis in BEAST 2.6.6 showed high convergence of all parameters in three independent runs after 25% of the trees were discarded as burn-in (ESS >300). The results were in good agreement with pervious results obtained from single genes (Alaei Kakhki et al. 2016), dating the origin of open-habitat chats to the Miocene about 5.67 million years ago (mya) (95% highest posterior density (HPD): 5.32-6.06 mya). The diversification of wheatears (genus *Oenanthe*) started about 5.09 mya (95% HPD: 4.75-5.44 mya) **(Fig. 1, Fig. S3).**

We then used the diversification time of the open-habitat chats estimated from mitochondrial data as a time constraint in dating analyses based on nuclear data. For these analyses, we first provided the topology and branch lengths obtained from ML analyses of concatenated BUSCO data along with 1.8 Mb high-confidence nuclear data (see Material and Methods) to generate the time-calibrated tree with RelTime-ML (Kumar et al. 2018). Compared to the mitochondrial results, the nuclear data mostly estimated similar divergence times between clades and shorter divergence times within clades (Pearson’s r=0.93, p<0.001) (**Fig. 1, Fig. S3**). Second, we performed dating analyses for windowed loci across the genome the same way as for BUSCO by providing 3.8 Mb high-confidence data. Divergence times based on BUSCO strongly correlated with ones estimated from windowed loci (Pearson’s r=0.99, p<0.001) **(Fig. S4)**. A test in which we re-ran the estimation of mitochondrial divergence times in RelTime-ML the same way as for nuclear data yielded the same divergence times as estimated in BEAST, thus confirming that differences in divergence times between mitochondrial and nuclear data are not due to the approach but reflect the different data types.

### Extensive Gene Tree Heterogeneity

Having established the species tree, we aimed to quantify the levels of gene tree heterogeneity in wheatears to understand whether the processes generating gene tree heterogeneity could underly convergent evolution in this core group of open-habitat chats that displays the highest incidence of convergent evolution.

Several lines of evidence demonstrate extensive gene tree heterogeneity in wheatears. Remarkably, not a single gene tree out of 29,730 gene trees matched the species tree. Furthermore, many branches of the species tree – including ones with local posterior probability 1 – showed a high number of conflicting compared to concordant bipartitions, as evidenced by low Internode Certainty All (ICA) scores (**Fig. S5**), with ICA ranging from 1 to 0.35 and average ICA of 0.65 ± 0.19 (mean ± standard deviation). The high gene tree heterogeneities highlighted by ICA were further supported by low percentages of gene trees recovering the topology of the species tree at these internodes, as estimated by the gene concordance factor (gCF) (**Fig. 1**) that ranged from 1 to 0.06 with an average of 0.52 ± 0.30 (mean ± standard deviation). ICA and gCF were highly correlated (Pearson’s r=0.94, p<0.001) **(Fig. S5)**. As expected, evidence for extensive gene tree heterogeneity was highest in clades with branches classified as within the phylogenetic anomaly zone. These included the *lugens, picata*, and *hispanica* complexes, the *isabellina* clade, and the placement of *O. leucopyga* and *O. leucura.*

### Contributions of ILS to Gene Tree Heterogeneity

Next, we aimed to understand to which extent the levels of gene tree heterogeneity observed in wheatears can be explained by ILS alone. To this end, we first tested whether the multi-species coalescent without hybridization adequately explains the gene tree heterogeneity observed across the entire species tree. The Tree Incongruence Checking in R (TICR) test (Stenz et al. 2015) showed an excess of outlier quartets (p < 0.01), indicating that a model including ILS but not introgression does not adequately explain the observed gene tree heterogeneity. This suggests that introgression occurred during the evolutionary history of wheatears.

Therefore, we moved on to infer for each branch in the species tree separately whether ILS alone may explain the level of gene tree heterogeneity. To this end, for each internal branch, we estimated the number of gene trees supporting the first and second alternative topologies, based on the rationale that under ILS the first and second alternative gene tree topologies should be supported by an equal number of gene trees (Sayyari and Mirarab 2018). We identified 11 out of 37 internal branches (30%) for which the number of gene trees supporting the two alternative topologies were not significantly different (colored branches in **Fig. 1**). At these 11 internal branches, ILS alone can thus explain gene tree heterogeneity, while asymmetries at the other 26 internal branches may need to invoke other processes.

### Contributions of Introgression to Gene Tree Heterogeneity

Given that gene tree heterogeneity at many branches could not be explained by ILS alone, we set out to infer footprints of introgression across wheatears. To this end, we first applied the approach based on D-statistics (Durand et al. 2011) implemented in Dsuite, using > 58 million biallelic SNPs. This approach estimates D and f4 statistics across all possible combinations of trios in wheatears and then performs an f-branch test to assign gene flow to specific internal branches. The f-branch test suggested multiple events of introgression (**Fig. 3**), namely between: (i) *O. halophila* and the ancestor of *O. lugens lugens* and *O. warriae*, (ii) *O. xanthoprymna* and the ancestor of the *lugens* complex, (iii) *O. leucopyga* and the ancestor of the *O. lugubris/lugentoides* clade, (iv) *O. picata capistrata* and the ancestor of *O. picata picata* and *O. albonigra*, and (v) *O. melanoleuca* and *O. pleschanka*.

**Figure 3.**
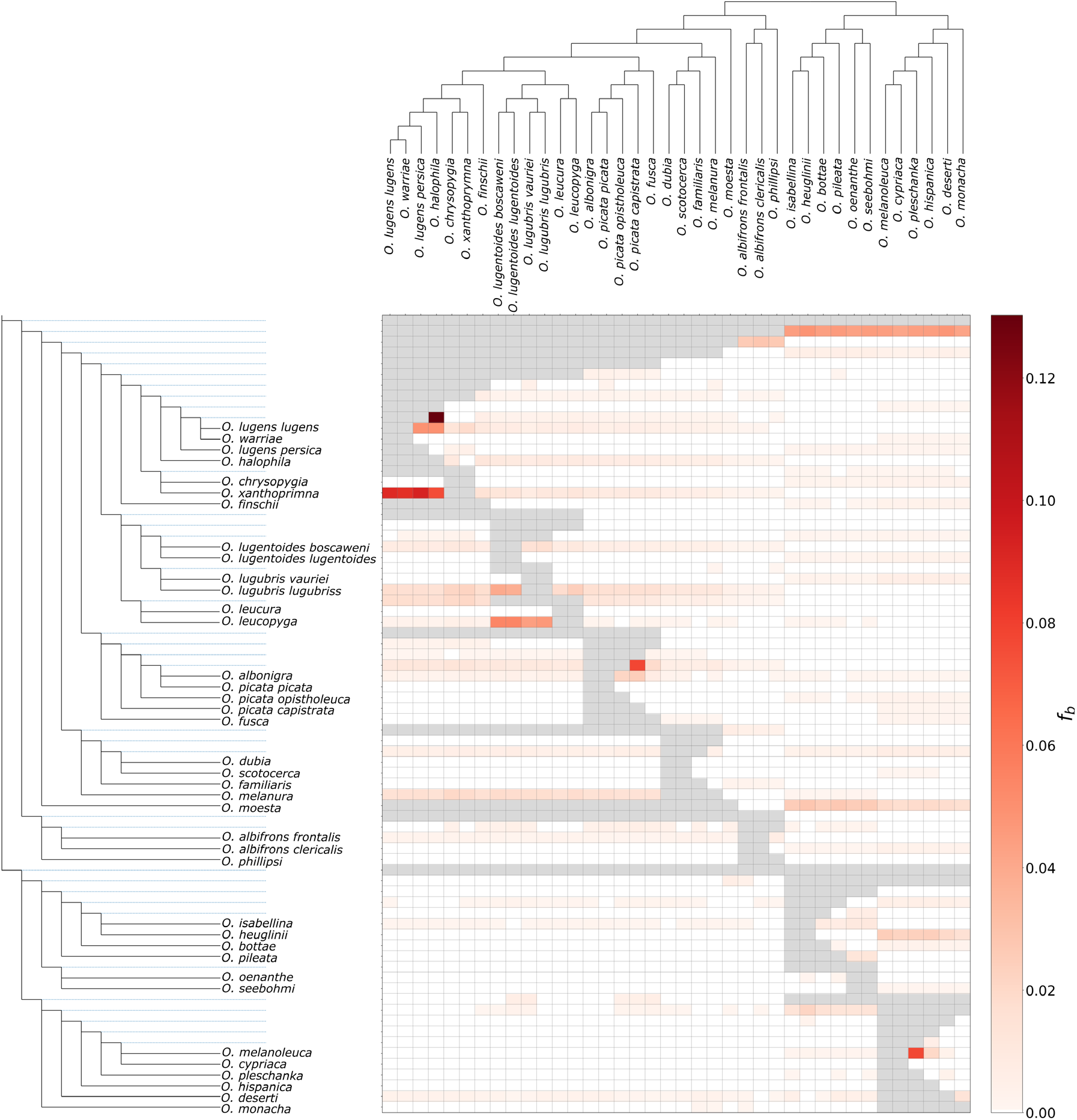
Footprints of introgression as estimated by the f-branch statistic. The heat map summarizes the f-branch statistics estimated in Dsuite. Darker colors depict increasing evidence for gene flow between lineages. Dotted lines in the phylogeny represent ancestral lineage.

Finally, we corroborated the evidence for introgression in the *hispanica, lugens*, and *picata* complexes with multi-species coalescent network analyses in phyloNet, allowing for 0-5 introgression events. According to the Bayesian Information Criterion (BIC), models involving reticulation events better fit the data than strictly bifurcating trees in all three complexes (**Tab. S3**). In the *lugens* complex, two introgression events were detected: between *O. xanthoprymna* and the ancestor of *O. lugens* (γ=49%), and between *O. halophila* and the *O. lugens lugens-O. warriae* ancestor (γ=25%) (**Fig. 4**). One introgression event was detected in the *picata* complex, between *O. picata capistrata* and the ancestor of *O. picata picata* and *O. albonigra* (γ=8%) (**Fig. 4**). In the *hispanica* complex, the highest-scoring network involved two introgression edges: one between *O. melanoleuca* and *O. pleschanka* (γ=17%), and one between *O. hispanica* and the *O. cypriaca-melanoeuca* ancestor (γ=1%) (**Fig. 4**).

**Figure 4.**
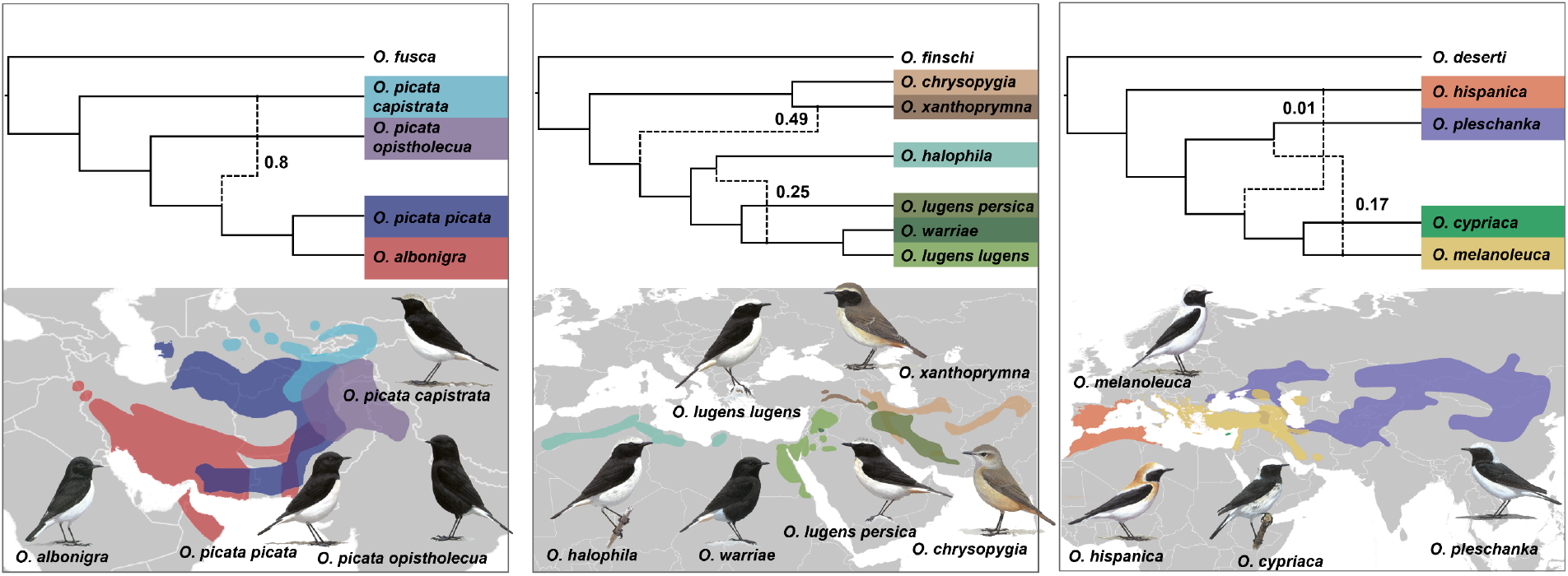
Phylogenomic networks and distribution ranges for the *picata* (left), *lugens* (middle) and *hispanica* (right) complexes. Phylogenomic networks were estimated under the maximum pseudolikelihood approach implemented in phyloNet. Numbers on the edges indicate the inheritance probabilities, which correspond to the proportion of gene trees supporting the reticulate relationship. Drawings courtesy of Chris Rose (www.chrisrose-artist.co.uk) with permission from Bloomsbury Publishing Plc. Distribution ranges modified from BirdLife International and the Handbook of the Birds of the World (2016)

## Discussion

The present study provides first genomic insights into the speciation history of open-habitat chats and into the processes involved in shaping gene tree heterogeneity that may also underpin the high incidence of convergent evolution in this group of songbirds. Our analyses reveal unambiguous species relationships despite considerable gene tree heterogeneity, including several mito-nuclear discordances that result from a combination of ILS and introgression. These relationships reconstructed from genomic data provide the strongest evidence yet for abundant convergent evolution in open-habitat chats, exemplified for three phenotypes in **Fig. 1**.

We first discuss how mito-nuclear discordances and incidences of introgression together with known histories of hybridization and biogeography mold into a comprehensive picture of open-habitat chat evolution. We close by concluding based on the indirect evidence presented here that convergent evolution in open habitat chats likely involved a combination of ILS, introgression and novel mutations in independent lineages. Together, our results paint a picture of genomic and phenotypic evolution that is in part marked by the sharing of ancestral variation and an exchange of genetic variation between species. Our study therefore contributes to the increasing body of evidence that phenotypic and species evolution not only proceed from novel mutations but abundantly reuse genetic variation present in ancestral and related species (Marques et al. 2019b; Meier et al. 2018; Seehausen et al. 2014).

### Mito-nuclear discordances, patterns of introgression, hybridization history, and biography mold into a coherent picture of complex open-habitat chat evolution

The species relationships inferred from nuclear genomic data were in good agreement with previous phylogenies based predominantly on single mitochondrial markers (Aliabadian et al. 2012; Schweizer and Shirihai 2013) and thereby confirmed the biogeographic history of open-habitat chats (Alaei Kakhki et al. 2016). Still, we recovered several species relationships discordant between the nuclear genome and the mitogenome (Toews and Brelsford 2012). In the light of (i) the histories of introgression also uncovered here, (ii) the previously known hybridization history deduced from observed instances of hybridization, and (iii) the here confirmed biogeography, most of these mito-nuclear discordances can be well embedded in a coherent history of open-habitat evolution.

The close nuclear relationship of *O. albonigra* with the nominate subspecies *O. p. picata* is in stark contrast with the mitochondrial divergence of *O. albonigra* with all *O. picata* subspecies about 0.5 mya (**Fig. 4a**). However, as an exception for wheatears, even from a perspective of plumage coloration, the nuclear species tree implies a more parsimonious history of phenotypic evolution, as *O. albonigra* and *O. p. picata* display almost identical plumages. The high mitochondrial similarity of all subspecies currently treated under *O. picata* according to Panov (2005) may be a result of introgressive hybridization. Indeed, the high abundance of admixed phenotypes in zones of contact between the members of this species complex (Panov 2005) suggests a high incidence of hybridization. Different from the *hispanica* complex, where taxa meet in restricted zones, lineages of the *picata* complex all mold together in a relatively large area in southern Central Asia, and their degree of reproductive isolation is largely unknown. Further population genomic insights are required from the *picata* complex to obtain detailed insights into its history of hybridization and phenotypic evolution.

The evolution of the *lugens* complex was marked by two incidences of introgression that likely underpin the mito-nuclear discordance observed in this complex (**Fig. 4b**). Introgression occurred between *O. xanthoprymna* and the *O. lugens* ancestor and between north-African *O. halophila* and the middle eastern *O. l. lugens-O. warriae* ancestor. Both incidences of introgression make sense in the light of biogeography, as they occurred between geographically neighboring taxa (**Fig. 4b**). Together they can explain the close mitochondrial relationship of *O. l. persica* with *O. xanthoprymna* and *O. chrysopygia: O. xanthoprymna* mitochondria were introduced into the *O. lugens* ancestor by hybridization and may at first have segregated in the *O. lugens* lineage but then have been lost in *O. halophila.* Mitochondrial replacement with *O. halophila* variation upon genetic exchange of the latter taxon with the *O. l. lugens-O. warriae* ancestor would have left *O. l. persica* the only taxon with a *O. xanthoprymna-*like mitogenome. Importantly, our results shed first genomic light on the divergence of Basalt Wheatear (*O. warriae*), a species with a very restricted range that is interesting from the perspective of phenotypic evolution: this species turns out to be highly similar to *O. l. lugens* at the genomic level, which contrasts with its marked phenotypic divergence (**Fig. 4b**). This result is similar to the situation observed, for instance, in Hooded and Carrion Crows (*Corvus cornix* and *C. corone*, respectively) (Poelstra et al. 2014) and opens interesting questions on the evolutionary history of this taxon’s coloration.

Finally, in the *hispanica* complex, the incomplete sorting of mitochondrial variation was previously well documented (Alaei Kakhki et al. 2018; Randler et al. 2012), and footprints of introgression came as no surprise: The complex is characterized by pervasive hybridization of *O. melanoleuca* with *O. pleschanka* in several geographic regions (Haffer 1977; Panov 1992) and population genomic analyses suggest rates of introgression of up to almost 20% between these species (Schweizer et al. 2019a). Research is underway to uncover the detailed histories of hybridization in this Eurasian wheatear complex.

The thus far discussed mito-nuclear discordances were all accompanied with high levels of gene tree heterogeneity (most within the phylogenetic anomaly zone). However, most of these cases were not explained by ILS alone but went along with footprints of introgression. Still, part of the observed mito-nuclear discordances might well be a consequence of ILS. In the *picata* complex, for instance, lineage divergence occurred in rapid succession (**Fig. 1**), and ILS might well be an alternative explanation for the mitochondrial divergence of the *O. albonigra* mitogenome. In addition, in the clade including *O. heuglinii* and the very widespread *O. isabellina*, species split in fast succession and the high levels of ILS likely explain the observed mito-nuclear discordance.

Taken together, our results demonstrate that the speciation history of open-habitat chats is similarly complex as their phenotypic evolution. Multiple events of introgression at both extant and ancestral time scales, along with abundant ILS, contributed to reticulate evolution and thus a mosaic of genomic variation in several clades of wheatears. Our study thus adds to an increasing number of examples (Enciso-Romero et al. 2017; Han et al. 2017; Lamichhaney et al. 2018; Meier et al. 2017) highlighting that species diversification is often complex and rather than by a linear process is at least in part a network of interacting lineages (Marques et al. 2019b)

### Diverse routes to convergent evolution in open-habitat chats

The reconstruction of relationships among open-habitat chats using genomic data has a deep impact on our understanding of phenotypic evolution in these songbirds: the species tree provides firm evidence for an extraordinary incidence of convergent evolution (**Fig. 1**). For numerous traits, including plumage coloration, sexual dimorphism, and migration behavior, not related species display more similar phenotypes than sister species (**Fig. 1**). Almost entirely black plumages, for instance, evolved in five clades (*O. picata opistholeuca*, *O. warriae*, *O. leucura*, female *M. monticola*, and juvenile *O. leucopyga*), and sexually monomorphic female-type plumage is found in another five clades (*O. chrysopygia, O. fusca*, the *O. melanura* clade, the *O. isabellina* clade, and in the sub-Saharan clade), to name just two out of many examples.

Furthermore, our results suggest (directly for introgression and ILS, indirectly for novel mutations) that convergent evolution in open-habitat chats is unlikely explained by a single process but may need to invoke all three processes (Hedrick 2013; Konečná et al. 2021; Montejo-Kovacevich et al. 2021; Natarajan et al. 2015; Pease et al. 2016), with the most likely processes depending on both demography and the phylogenetic scale.

For ILS to contribute to convergent evolution, species must diverge in fast succession and maintain critically high effective population sizes to pass on ancestral variation and maintain it in daughter lineages. In open-habitat chats, such fast radiations occurred predominantly at rather recent time scales. The shortest split intervals are observed (in increasing order) in the *picata*, *hispanica*, and *lugens* complexes (**Fig. 1**). However, convergent evolution of species in the *lugens* complex and of the *picata* complex is only found with other clades but not within the complexes. Given that the levels of ILS at the root of the *lugens* complex are restricted, ILS is unlikely to have contributed to convergent evolution with other clades of wheatears sporting, for instance, similar plumages (see for instance the aforementioned example including *O. warriae).* Convergent evolution is, however, observed for back and neck-side coloration in the *hispanica* complex (Schweizer et al. 2019a), and could be explained by ILS of ancestral variation.

Likewise, introgression would need to have happened between taxa with similar phenotype to explain convergent evolution. Our analyses indeed uncovered several instances of in part substantial introgression (**Fig. 3, Fig. 4**). However, despite suggesting that introgression upon hybridization provided the opportunity to exchange phenotypes between species, none of the inferred introgression events can be tied to concrete examples of convergent evolution. This raises the question, whether the methods applied here are underpowered to infer footprints of introgression relevant to phenotypic evolution of open-habitat chats, or, indeed, introgression played a limited role in these songbirds’ phenotypic evolution.

Finally, we may need to invoke novel mutations to explain at least part of the observed convergent evolution, because phenotypic similarities are found between rather divergent species and inferred instances of high ILS and introgression cannot easily explain them. Many if not most phenotypic similarities in open-habitat chats are found in the rather distant major phylogenetic clades that diverged around 5 mya (for instance the examples provided at the entry of the discussion). The time tree suggests that the relevant split events did not occur within short evolutionary time scales. Accordingly, levels of ILS are rather low for at least one of the relevant nodes (**Fig. 1**). Although gene tree heterogeneity was non-negligible for the larger of the two major wheatear clades, gene trees were mostly concordant for the root nodes of the wheatear clade including the *hispanica* complex and the *O. oenanthe* and *O. isabellina* clades (**Fig. 1**). Moreover, the phenotypically similar species occur in geographically well separated ranges and introgression between them is thus rather unexpected. In conclusion, unless the approaches used here to detect the ILS and introgression are underpowered, the indirect evidence provided by our results suggests that many incidences of convergent evolution at such time scale may have involved independent novel mutations.

### Conclusion

In the present study we set out to probe gene tree variation for footprints of ILS and introgression with the goal of understanding how ILS and introgression may have contributed to convergent evolution in open-habitat chats. Our results reveal a complex speciation history and provide conclusive evidence for abundant convergent evolutionin open-habitat chats. While we cannot conclude on the involvement of specific processes in the evolution of specific convergent evolution, the indirect evidence gained from the structure of the species tree and inferred levels of ILS and introgression suggest that convergent evolution in open-habitat chats likely occurred via all three possible processes, namely ILS, introgression, and novel mutations. Thereby, our results contribute to a growing body of evidence that evolution makes use and re-use of all resources it has at hand, including both standing (ancestral or heterospecific) as well as novel genetic variation.

Finally, the approach applied here based predominantly on a characterization of gene tree heterogeneity outlines an avenue to probe the processes governing convergent evolution in a wide range of systems. Even though the evidence for the involvement of these processes is indirect, ultimately, at a comparative scale this evidence may provide valuable insights into the relative contributions of ILS, introgression, and novel mutations to convergent evolution.

## Material and Methods

### Taxon sampling, DNA extraction, and whole-genome resequencing

Aiming for complete taxon sampling, we sequenced the genomes of 50 open-habitat chat taxa from a total of 44 species from the genera *Oenanthe, Campicoloides, Emarginata, Myrmecocichla, Pinarochroa*, and *Thamnolaea* (**Fig. 2**; **Tab. S1**). This sampling included all but three species (*E. tractrac, M. collaris, T. coronata*) of the 47 currently recognized open-habitat chat species (Gill et al. 2020). A genome sequence of *Saxicola maurus* (European Nucleotide Archive accession number: ERR2560200-ERR2560209), a species of open-habitat chats’ sister lineage (Sangster et al. 2010; Zuccon and Ericson 2010), was included as an outgroup to root the open-habitat chat species tree. We followed the taxonomy of the IOC World Bird List (v12.1) (Gill et al. 2020) except for the *picata* complex, where we treat subspecies *picata, capistrata* and *opistholeuca* separately, following Panov (2005).

We extracted DNA from blood stored in ≥96% ethanol or Queen’s Lysis buffer or tissues stored in 96% ethanol for taxa for which fresh material was available, or from toepads or dried skin from skin-preparation sutures for taxa for which only museum samples were available (**Tab. S1**). From blood and tissue samples DNA was extracted using the DNeasy Blood and Tissue Kit (Qiagen) or the MagAttract HMW DNA kit (Qiagen) following the manufacturer’s protocol with exception of an adapted digestion of blood samples as reported in Lutgen et al (Lutgen et al. 2020). DNA from toepads and dried skin was extracted using the QIAamp DNA Micro Kit (Qiagen) with an adapted digestion protocol that ensures high quantities of DNA (dx.doi.org/10.17504/protocols.io.dm6gpwrdplzp/1). DNA concentrations were quantified on a Qubit fluorometer (dsDNA BR assay, Thermo Fisher Scientific) and DNA integrity was evaluated on a TapeStation (MANUFACTURER, KIT). We prepared sequencing libraries using the ThruPLEX DNA-Seq Kit (Takara), the Illumina DNA Prep Kit, the Illumina DNA PCR-free Kit, or the Chromium Genome Library kit (10X Genomics) for intact DNA, or for fragmented DNA with the ACCEL-NGS 1S DNA Library Prep Kit (Swift Biosciences) (**Tab. S1**). All libraries were sequenced (150 bp paired-end) on Illumina NovaSeq6000 instruments with a target coverage of ca. 15x.

### Data preparation

#### Adapter trimming and mapping of resequencing data

Prior to further analysis, for all but the linked-read sequencing data, we trimmed adapters and merged overlapping paired-end reads using fastp 0.20.0 (Chen et al. 2018). For linked-read sequences, we trimmed the first 22 bp on the R1 read to eliminate the 10X indexes. We then mapped the reads to the reference genome assembly of *Oenanthe melanoleuca* (Peona et al. in prep.) using BWA 0.7.17 (Li 2013) and marked duplications with PicardTools 2.9.1 (http://broadinstitute.github.io/picard). After excluding duplicates, the average sequencing coverage per individual ranged from 4.6x to 40.6x (mean and median 12.2, standard deviation 6.20) (**Tab. S1**).

#### Base quality score recalibration (BQSR), SNP calling, and SNP genotyping

Data preparation followed the GATK 4.1.4.1 (McKenna et al. 2010) best practices pipeline. First, to prepare a list of high-confidence SNPs for BQSR, we ran HaplotypeCaller to generate gvcf files for each sample and then merged gvcf files of all samples with CombineGVCFs before genotyping SNPs using GenotypeGVCFs. To retain only high-confidence SNPs in the SNP-exclude set for BQSR, we retained only SNPs that fulfilled the following criteria: mapping quality > 40, Fisher strand (FS) phred-scaled p-value < 60, SNP quality score > 20, mapping quality rank sum value > −12.5, read pos rank-sum test value > −8.0 and quality by depth > 2. We retained only biallelic SNPs with at least one homozygous reference and one homozygous alternative genotype or with at least three observations of reference and alternative alleles. We excluded the resulting set of SNPs from BQSR in GATK. Following BQSR, we ran HaplotypeCaller on base-score-recalibrated bam files. The resulting gvcf files of all samples where merged (CombineGVCFs) and variant and invariant sites genotyped using the ‘include-non-variant-sites’ flag in GenotypeGVCFs. For all subsequent analyses we based genotypes on genotype likelihoods. This resulted in 871,428,254 unfiltered sites when the outgroup was included and 872,152,150 unfiltered sites without the outgroup.

In phylogenomic data sets, which are based on mapping of resequencing data to a reference genome, data of species more divergent from the reference genome may risk mapping at a lower percentage. To check for such mapping-related biases in our dataset, we estimated the average number of nucleotide differences (d_*XY*_) between *Oenanthe melanoleuca* (reference genome) and all other species using pixy 0.95.02 (Korunes and Samuk 2021). We then estimated the mapping percentage for all species using SAMtools (Li et al. 2009) and tested whether there was a correlation between d_*XY*_ and mapping success.

#### Data filtering

Before data analysis, we removed all repeat regions from the multi-sample VCF file using the repeat mask reported in Peona et al. (in prep.). Then we used BCFtools 1.11 (Li 2011) to remove indels, sites close to indels (up to 10 bp) and all the sites at which exclusively alternative alleles were called. For analyses requiring variant sites only, we removed all SNPs with more than 20% missing data and all invariant sites using BCFtools and retained only SNPs with a minimum read depth of five. To ensure linkage-disequilibrium (LD) among SNPs, we LD-pruned SNPs in VCFtools 0.1.16 (Danecek et al. 2011) such as to only retain SNPs with a minimum distance of 1 kb between them. This physical distance is expected to remove most LD between SNPs, as e.g. in flycatchers LD breaks down in most genomic regions after 1 kb (Ellegren et al. 2012). After this filtering, we genotyped based on genotype likelihoods and retained 994,150 multiallelic SNPs. In addition, for analyses that require biallelic SNPs exclusively, we removed all multiallelic SNPs from the VCF file after the above filtering, using BCFtools.

For phylogenomic analyses requiring sequence data including both variant and invariant sites, we followed two strategies. First, we defined 10 kb non-overlapping windows across the genome. Henceforth, we refer to the windowed data as “loci” and to phylogenetic trees inferred therefrom as “gene trees”. Second, we inferred benchmarking universal single-copy orthologs, BUSCO, using BUSCO 5.0.0 (Simão et al. 2015). Similar to ultraconserved elements, UCE (Faircloth et al. 2012), BUSCO feature a high degree of conservation and moreover are present in single copies, circumventing issues with paralogs in phylogenomic reconstructions (Roy 2009). BUSCO are readily identified in whole-genome resequencing data sets, not requiring genome alignments, and are increasingly deployed for phylogenomic reconstructions (Kallal et al. 2021; Van Damme et al. 2022).

To make sure that the adopted filtering strategy did not affect our results, we generated four sets of fasta alignments using different filter settings for minimum read depth (DP), minimum percentage of the window covered by data (PW), and missing data per site (MD) for both the 10 kb loci and the BUSCO data set: (i) DP=1, PW=50%, MD=15%, (ii) DP=5, PW=50%, MD=15%, (iii) DP=1, PW=50%, MD=5%, and (iv) DP=1, PW=80%, MD=10% (**Tab. S2**). These four filtering strategies yielded the same species tree and concatenated tree for 10 kb loci as well as for BUSCO. For these analyses, we therefore exclusively report the results based on the most stringent filtering on read depth (ii, DP=5, PW=50%, MD=15%) (**Tab. S2**). For gene tree heterogeneity analyses, on the other hand, we aimed to include the broadest representation of the genome and to this end retained all loci (N=29,730) that fulfilled less stringent filtering criteria (i, DP=1, PW=50%, MD=15%) (**Tab. S2**).

Finally, for analyses making assumptions on intra- and inter-locus recombination (such as species tree reconstructions) we made sure to include only loci with no intra-locus but free inter-locus recombination **(Tab. S2**). To this end, we excluded all loci with recombination signals (P ≤ 0.05) as inferred from the pairwise homoplasy index Phi (Φw) estimated in PhiPack 1.1 program (Bruen et al. 2006). The criterion P ≤ 0.05 does not account for multiple testing, but we preferred to conservatively exclude loci with evidence for intra-locus recombination. To possibly retain only loci among which free recombination occurs, we ensured a minimum distance of 10 kb by including no two consecutive loci. At this distance, no LD occurs in flycatchers (Ellegren et al. 2012).

#### Inference of Benchmarking Universal Single-Copy Ortholog (BUSCO) sequences

Phylogenomic analyses based on the mapping of resequencing data to a reference genome, especially when including species well diverged from the latter, may be affected by several biases. For species more divergent from the reference genome, data from faster evolving genomic regions (i) risks not being mapped, if these regions are too diverged from the reference sequence, or (ii) may map to paralogs, if the species experienced different duplication histories (Chakrabarty et al. 2017; Fitz-Gibbon et al. 2017). These biases are expected to be least important in slowly evolving regions of the genome, especially in BUSCO, that are conserved and by definition present in single copies in most species. To minimize mapping-related biases in our phylogenomic reconstructions, especially on rooting and placements of the most divergent species, we therefore extracted the intervals in which avian BUSCO (aves_odb10) are situated in our reference genome using BUSCO 5.0.0 (Simão et al. 2015).

### Phylogenomic reconstructions and multispecies coalescent analyses

#### BUSCO-based rooting of the open-habitat chat species tree

First, to establish the root within open-habitat chats, we applied both concatenation and multispecies coalescent-based methods on BUSCO sequences, including the outgroup. First, we used all BUSCO (N=7,335) to estimate the maximum likelihood tree in IQ-TREE 2.1.2 (Minh et al. 2020b) based on the concatenated BUSCO, using with one partition for each BUSCO and a GTR+I+G substitution model for all partitions (Abadi et al. 2019). One thousand bootstrap replicates were run using the ultrafast bootstrap approximation (Hoang et al. 2018). Second, we estimated the species tree under the multispecies coalescent using ASTRAL-III (Zhang et al. 2018) based on BUSCO without recombination signals and free inter-locus recombination (N=2,091). To this end, we inferred BUSCO’ gene trees in IQ-TREE 2.1.2 using a GTR+I+G substitution model and one thousand ultrafast bootstrap approximations. To ensure that species tree inferences were not affected by inaccurately estimated gene trees (Zhang et al. 2018), we collapsed branches with bootstrap support inferior to 80% using Newick Utilities 1.6 (Junier and Zdobnov 2010). Reconstructing the species tree by including all BUSCO not considering intra- and inter-locus recombination (N=7,335) did not affect the result.

#### Phylogenomic and multispecies coalescent analyses based on full evidence

To reconstruct the concatenated tree and species tree based on full evidence data, that is, data from the maximal possible fraction of the genome, and to study gene tree heterogeneity along the genome, we excluded the *Saxicola* outgroup. Instead, we rooted the trees with the clade that is the outgroup to all other open-habitat chats (sub-Saharan clade, **Fig. S1**). Excluding *Saxicola* ensured that analyses were not biased by mapping issues caused by this outgroup’s divergence.

To estimate the concatenated tree using maximum likelihood in IQ-TREE 2.1.2 we used all loci with a GTR+I+G substitution model and 1,000 ultrafast bootstrap approximations. To estimate the species tree under the multispecies coalescent using ASTRAL-III, we at first estimated maximum likelihood gene trees using IQ-TREE 2.1.2 with a GTR+I+G substitution model and one thousand ultrafast bootstrap approximations. Based on these gene trees (pruned for within-locus recombination and assuring free recombination between loci), we inferred the species tree using ASTRAL-III. Because ASTRAL relies on accurately estimated gene trees, we collapsed branches with bootstrap support inferior to 80% using Newick Utilities 1.6.

To find regions of the species tree that represent “anomaly zones” where the frequency of one of the alternative quartets is higher than that of the topology in agreement the species tree, we estimated local quartet supports for the main topology and its two alternatives in ASTRAL-III (Degnan and Rosenberg 2006). We used the anomaly_finder.py script to search for anomaly zones in our species tree (Linkem et al. 2016). To test if the gene tree discordance could be explained by polytomies instead of bifurcating nodes, we carried out a quartet-based polytomy test as implemented in ASTRAL-III.

To see whether the SNP-based species tree could confirm the sequence-based species tree, we used the unlinked multiallelic SNPs to the multispecies coalescent model implemented in SVDQuartets (Chifman and Kubatko 2014) in PAUP* 4 (Swofford 2003). We ran this with 1000 bootstrap replicates and summarized the result in a 50% majority-rule consensus tree.

#### Phylogenetic relationships of mitogenomes

We were interested in whether previously inferred relationships based predominantly on single mitochondrial genes (Alaei Kakhki et al. 2016; Aliabadian et al. 2012; Schweizer and Shirihai 2013) were supported by full mitogenomes and in how the mitogenomic relationships compare to the ones inferred from nuclear loci. To this end, we extracted and assembled mitochondrial genomes from the genomic data of all open-habitat chats using MitoFinder 1.2 (Allio et al. 2020). We used the published Isabelline Wheatear (*Oenanthe isabellina*) mitochondrial genome as a reference (Genbank accession number: NC_040290.1) and annotated the mitochondrial genome using the annotation pipeline integrated in MitoFinder. Finally, we aligned the 13 mitochondrial protein coding gene sequences using the automatic alignment strategy in MAFFT 7.471 (Katoh and Standley 2013). We checked the alignments in AliView 1.26 (Larsson 2014) and removed stop codons within the coding sequences or indels for downstream analyses. We determined the best partition scheme using the Akaike information criterion (AIC) implemented in PartitionFinder 2.1.1 (Lanfear et al. 2017) and used the GTR+G+I model for all partitions. Then we constructed the maximum-likelihood tree from the concatenated supermatrix of all 13 genes in IQ-TREE 2.1.2 using the ultrafast bootstrap approximations with 1,000 replicates.

### Dating analyses

Beside species’ relationships we were interested in estimating the divergence time in open-habitat chats. Because there are no appropriate fossils for calibration, we first ran BEAST 2.6.6 (Bouckaert et al. 2019) for 13 mitochondrial protein coding genes to estimate a time-calibrated mitochondrial phylogeny. We included the mitochondrial genome sequence of *Saxicola maurus* (GenBank accession number: MN356403.1) as an outgroup in these analyses. Substitution models were inferred during the MCMS analyses with bModelTest (Bouckaert and Drummond 2017) implemented as a package in BEAST 2.6.6. Published substitution rates for each mitochondrial gene (Lerner et al. 2011) were implemented as means of the clock rates in real space of lognormal distribution with standard deviations of 0.005. We defined a Yule speciation process for the tree prior and an uncorrelated lognormal relaxed clock model. Three independent MCMC chains were run for 50 million generations, each with sampling every 5,000 generations. Effective sample sizes for all parameters and appropriate numbers of burn-in generations were checked with Tracer 1.5 (Rambaut and Drummond 2009). The three independent runs were combined using LogCombiner 2.6.6 (Bouckaert et al. 2019). We used TreeAnnotator 2.6.6 (Bouckaert et al. 2019) to calculate a maximum clade credibility tree and the 95% highest posterior density (HPD) distributions of each estimated node.

We then used the divergence time of the sub-Saharan clade from wheatears estimated from mitochondrial data as time constraint in dating analyses based on nuclear data using RelTime-ML implemented in MEGA 11 (Tamura et al. 2021). For this analysis, we provided the topology with branch length estimated in IQtree2 based on concatenated BUSCO data retained after the most stringent filtering (ii, DP=5, PW=50%, MD=15%), along with high-confidence BUSCO alignments. The latter consisted of BUSCO data filtered for DP=5, MD=5% and length of each BUSCO alignments longer than 1kb. We used the same filtering to get the 10 kb non-overlapping windows across the genome and used the concatenated tree retained after most stringent filtering (ii, DP=5, PW=50%, MD=15%) to repeat the analyses based on loci across the genome. To ensure that the differences in divergence times between mitochondrial and nuclear data were not due to the different dating approaches, we re-estimated the mitochondrial divergence times in RelTime-ML using the same approach as for the nuclear datasets.

### Inference of gene tree variation, ILS, and introgression

#### Inference of the levels of gene tree variation

To investigate gene tree heterogeneity across the genome, we used gene trees inferred from less stringent filtering criteria (i, DP=1, PW=50%, MD=15%) as described above. To infer how many gene trees reflect the species tree topology, we used the script ‘findCommonTrees.py’ (Edelman et al. 2019). To characterize the levels of gene tree heterogeneity across open-habitat chats, we compared the gene trees to the species tree. Specifically, we estimated “internode certainty all” (ICA) and the “gene concordance factor” (gCF). ICA quantifies the amount of gene tree heterogeneity for each internode of the species tree by calculating the number of all most prevalent conflicting bipartitions. It takes values ranging from −1 to 1, with values around zero indicating strong conflict; values towards 1 indicate robust concordance of gene trees with the species tree in the bipartition of interest; and negative values indicate discordance between the bipartition of interest and one or more bipartitions with a higher frequency (Salichos et al. 2014). While ICA thus represents the degree of conflict on each node of a species tree, gCF better reflects the gene tree heterogeneity around each branch, and is the percentage of gene trees supporting the two alternative topologies for each branch (Minh et al. 2020a). We estimated ICA and gCF with PhyParts 0.0.1 (Smith et al. 2015) and IQ-TREE 2.1.2 respectively.

#### Tests of an ILS model

Next, we were interested in understanding whether ILS can sufficiently explain the level of gene tree heterogeneity observed at the level of the whole species tree. To this end, we applied the Tree Incongruence Checking in R (TICR) test (Stenz et al. 2015) implemented in the Phylolm R package. This test evaluates whether the multispecies coalescent adequately explains gene tree heterogeneity across the species tree with no hybridization edges. TICR requires posterior distributions of gene tree topologies inferred through Bayesian inference of gene trees. Therefore, we first estimated posterior distributions of individual gene trees with MrBayes 3.2.7 (Ronquist et al. 2012). MrBayes analyses ran using three independent runs of 20 million generations each, sampling every 20,000th generation using a GTR+I+G model. We estimated the length of burn-in using Tracer 1.5 (Rambaut and Drummond 2009) to ensure that our sampling of the posterior distribution had reached sufficient effective sample sizes (ESS > 200) for parameter estimation. We then ran BUCKy (Ané et al. 2007; Larget et al. 2010) using the posterior distribution of gene trees after discarding 25% as burn-in to estimate the concordance factors (CFs) for the three possible splits of all quartets. The inferred CF values were then tested against those expected under a coalescent model that takes ILS but not hybridization into account (chi-squared test).

We then tested for each branch in the species tree whether the gene tree heterogeneity reflected in gCF can be sufficiently explained by a model incorporating ILS alone. Under ILS alone – assuming sorting of variation occurs by random genetic drift – proportions of alternative gene trees for a rooted triplet are expected to be approximately equal (Hibbins and Hahn 2022; Sayyari and Mirarab 2018; Sayyari et al. 2018), and the concordant tree topology (the topology in agreement with the species tree) should be at least as frequent as the two discordant topologies (Hibbins and Hahn 2022; Sayyari et al. 2018). In contrast, introgression between non-sister taxa results in asymmetric proportions of gene trees in the rooted triplet (Durand et al. 2012; Green et al. 2010). Therefore, we performed a chi-square tests comparing the number of gene trees supporting the two discordant topologies. Under ILS, these two alternative topologies are expected to be equally frequent among gene trees (He et al. 2020). For all these analyses we accounted for uncertainty in gene tree topologies by collapsing branches with bootstrap support <80%.

#### Inferring footprints of introgression

To infer footprints of introgression across the entire species tree, we estimated Patterson’s D (Durand et al. 2011) and related statistics in Dsuite (Malinsky et al. 2021) based on 58,963,109 biallelic SNPs. D and f4 statistics were estimated across all possible combinations of trios in our 38 wheatear taxa. We used Dtrios to calculate the sums of three different patterns (BABA, BBAA and, ABBA) and D and f4-ratio statistics for all 8,437 possible trios. Dsuite uses the standard block-jackknife procedure to assess the significance of the D statistic. Due to the large number of D-statistics comparisons and difficulties disentangling false positives that may arise due to ancient gene flow, we performed the f-branch test (fb) implemented in Dsuite to assign gene flow to specific internal branches on the species tree. Then we visualized the output using Dsuite’s dtools.py script.

We then aimed to obtain further support for the footprints of introgression that were suggested in *lugens, picata* and *hispanica* complex by the above approach based on the D-statistics. To this end, for these three complexes, we estimated phylogenetic networks from maximum likelihood trees generated from BUSCO using the pseudolikelihood (InferNetwork_MPL) (Yu and Nakhleh 2015) and likelihood (CalGTProb) (Yu et al. 2014) approaches implemented in phyloNet 3.6.9 (Than et al. 2008). Due to the high computational demands, analyses were run for each of the clades containing signals of introgression in earlier analyses separately, namely for the *lugens, picata* and *hispanica* complexes. Furthermore, we only included BUSCO loci that had data available for all taxa of the respective complex. Outgroup species for each complex were selected based on the species tree. Analyses included 7,323 rooted gene trees for the *lugens* complex, 7,310 rooted gene trees for the *picata* complex, and 7,335 rooted gene trees for the *hispanica* complex. For each complex, we allowed for one to five reticulation events, with the starting tree corresponding to the species tree topology (-s), 0.9 bootstrap threshold for gene trees (-b) and 1,000 iterations (-x). To ensure convergence, the network searches were repeated 10 times. Then we estimated the likelihood by fixing the topology of the focal clade for the species tree (without any reticulation) and for each of the five networks (with different numbers of introgression edges) and calculated their likelihood scores. We determined the optimal network by calculating the Bayesian Information Criterion (BIC) from the maximum likelihood scores, the number of gene trees, the number of branch length being estimated, plus the number of admixture edges in each model (**Tab. S3**). We used the browser-based tree viewer IcyTree (Vaughan 2017) to visualize the estimated networks.

## Acknowledgements

We thank all the natural history museums and their staff who provided material for this study: namely the American Museum of Natural History, New York; Natural History Museum, Tring; Field Museum of Natural History, Chicago; Natural History Museum of Los Angeles County, Los Angeles; Museum of Vertebrate Zoology, UC Berkeley; Natural History Museum, University of Oslo; Naturhistoriska Riksmuseet, Stockholm; Texas A&M University Biodiversity Research and Teaching Collections, Texas College Station; University of Washington Burke Museum;, Seattle, Yale Peabody Museum; Zoological Museum, Natural History Museum of Denmark, Copenhagen; Zoologisches Forschungsmuseum König, Bonn; and Martin Haase, Vogelwarte Hiddensee, Universität Greifswald. Further samples were provided by José Luis Copete, and Marc Illa. We are indebted to the sequencing facilities of NGI Sweden in Solna and to the NGS platform of the University of Berne and their respective staff for their excellent services and to Marta Burri for sequencing library preparations. Computations were performed at the High-Performance Computing Cluster EVE, a joint effort of the Helmholtz Centre for Environmental Research (UFZ) and the German Centre for Integrative Biodiversity Research (iDiv) Halle-Jena-Leipzig. We thank the administration and support staff of EVE, Thomas Schnicke and Ben Langenberg (UFZ), and Christian Krause (iDiv). We thank Chris Rose and Claire Weatherhead from Bloomsbury Publishing Plc for their permission to use bird drawings in our figures. This research was supported by a German Research Foundation (DFG) research grant (BU3456/3-1) to RB, the National Research Fund (FNR), Luxembourg, grant number 14575729 to DL, and a Georg Foster Research Stipend of the Alexander von Humboldt Foundation and a scholarship for female researchers from Friedrich-Schiller-University Jena, both to NAK.

## Author Contributions

RB and NAK designed the study. NAK, DL and MS performed data analysis with inputs from HSch and RB. RCKB, AS and HShi provided materials. MS designed the figures. NAK and RB wrote the manuscript with help from MS and HSch and inputs from all authors.

## Data Availability

All sequencing data produced in the framework of this study are available on the European Nucleotide Archive (ENA) under accession number *XYZ (to be announced upon acceptance of the article*).

