## Supplementary Material for "A Phylogenomic Assessment of Processes Underpinning Convergent Evolution in Open-Habitat Chats"

### Supplementary tables

**Table S1 | Sampling and sequence data information.**

| Taxon | Collection, no.[a] | Tissue | Library Type | Mapping % | Coverage |
|---|---|---|---|---|---|
| *Campicoloides bifasciatus* | MVZ, RSA073, | Muscle | Illumina DNA Prep Kit | 99.1 | 8.5 |
| *Emarginata schlegelii* | FMNH, 453197, | Muscle | Illumina DNA Prep Kit | 99.1 | 9.5 |
| *Emarginata sinuata* | UWMB, 95470 | Muscle | ThruPLEX DNA-Seq Kit | 96.7 | 9.7 |
| *Myrmecocichla aethiops* | MVZ, 129113 | Toepad | ACCEL-NGS 1S DNA Library Prep Kit | 93.3 | 6.8 |
| *Myrmecocichla arnotti* | FMNH, 468111 | Toepad | Illumina DNA Prep Kit | 99.3 | 12.8 |
| *Myrmecocichla formicivora* | MVZ, RSA205 | Muscle | Illumina DNA Prep Kit | 98.6 | 8.3 |
| *Myrmecocichla melaena* | A1153 | Blood | ThruPLEX DNA-Seq Kit | 98.8 | 13.3 |
| *Myrmecocichla monticola* | NMBE, 1043860 | Toepad | ACCEL-NGS 1S DNA Library Prep Kit | 94.3 | 7.6 |
| *Myrmecocichla nigra* | NRM, 570041 | Toepad | ThruPLEX DNA-Seq Kit | 94.5 | 7.0 |
| *Myrmecocichla tholloni* | YPM, ORN95640 | Toepad | ThruPLEX DNA-Seq Kit | 92.1 | 7.3 |
| *Oenanthe (C.) dubia* | FMNH, 83201 | Toepad | ACCEL-NGS 1S DNA Library Prep Kit | 93.0 | 6.8 |
| *Oenanthe (C.) familiaris* | MVZ, GO866 | Blood | Illumina DNA Prep Kit | 99.4 | 13.3 |
| *Oenanthe (C.) fusca* | YPM, ORN011707 | Toepad | ThruPLEX DNA-Seq Kit | 86.4 | 4.7 |
| *Oenanthe (C.) melanura* | A1203 | Blood | ThruPLEX DNA-Seq Kit | 98.8 | 15.4 |
| *Oenanthe (C.) scotocerca* | LACM, 61131 | Toepad | ACCEL-NGS 1S DNA Library Prep Kit | 96.6 | 8.3 |
| *Oenanthe (M.) albifrons clericalis* | NRM, 558941 | Toepad | ThruPLEX DNA-Seq Kit | 95.4 | 6.3 |
| *Oenanthe (M.) albifrons frontalis* | KU, 115365 | Muscle | ThruPLEX DNA-Seq Kit | 98.5 | 12.7 |
| *Oenanthe albonigra* | IR-KIL-010 | Blood | ThruPLEX DNA-Seq Kit | 99.2 | 12.9 |

| | | | | | |
|---|---|---|---|---|---|
| *Oenanthe bottae frenata* | NRM, 558917 | Toepad | ThruPLEX DNA-Seq Kit | 97.0 | 6.0 |
| *Oenanthe chrysopygia* | IR-FIR-002 | Blood | ThruPLEX DNA-Seq Kit | 98.9 | 11.5 |
| *Oenanthe cypriaca* | 19e | Blood | Chromium Genome Library kit | 99.8 | 40.6 |
| *Oenanthe deserti* | MO-BOULMANE-2013 | Blood | ThruPLEX DNA-Seq Kit | 99.3 | 12.6 |
| *Oenanthe finschii* | IR-ESF-004 | Blood | ThruPLEX DNA-Seq Kit | 98.9 | 8.7 |
| *Oenanthe halophila* | 3Y42902 | Blood | Illumina DNA PCR-free | 99.6 | 17.4 |
| *Oenanthe heuglinii* | ZFMK, H.II.16p2.α | Dry skin | ACCEL-NGS 1S DNA Library Prep Kit | 93.6 | 5.7 |
| *Oenanthe hispanica* | E-GUI-013 | Blood | Chromium Genome Library kit | 99.8 | 15.5 |
| *Oenanthe isabellina* | GR-LES-001 | Blood | ThruPLEX DNA-Seq Kit | 99.2 | 9.9 |
| *Oenanthe leucopyga leucopyga* | A1137 | Blood | ThruPLEX DNA-Seq Kit | 98.8 | 10.7 |
| *Oenanthe leucura leucura* | E-MAT-2012 | Blood | ThruPLEX DNA-Seq Kit | 99.0 | 9.5 |
| *Oenanthe lugens lugens* | 9b | Blood | ThruPLEX DNA-Seq Kit | 98.7 | 22.0 |
| *Oenanthe lugens persica* | ZMUC, 137759 | Muscle | ThruPLEX DNA-Seq Kit | 98.6 | 10.4 |
| *Oenanthe lugentoides lugentoides* | NHMUK, 1965.M.12140 | Toepad | ThruPLEX DNA-Seq Kit | 96.9 | 4.6 |
| *Oenanthe lugentoides boscaweni* | NHMUK, 1977.M.21.36 | Toepad | ThruPLEX DNA-Seq Kit | 97.2 | 6.8 |
| *Oenanthe lugubris lugubris* | A1129 | Blood | ThruPLEX DNA-Seq Kit | 99.0 | 12.5 |
| *Oenanthe lugubris vauriei* | AMNH, 461151 | Toepad | ThruPLEX DNA-Seq Kit | 95.9 | 5.3 |
| *Oenanthe melanoleuca* | IT-GRA-006 | Blood | Chromium Genome Library kit | 99.7 | 12.6 |
| *Oenanthe moesta* | A1109 | Blood | ThruPLEX DNA-Seq Kit | 98.8 | 12.0 |
| *Oenanthe monacha* | A1174 | Blood | ThruPLEX DNA-Seq Kit | 99.3 | 17.2 |
| *Oenanthe oenanthe* | GEO-VAR-002 | Blood | ThruPLEX DNA-Seq Kit | 99.1 | 12.2 |

| | | | | | |
|---|---|---|---|---|---|
| *Oenanthe phillipsi* | YPM, ORN035210 | Toepad | ThruPLEX DNA-Seq Kit | 95.6 | 7.9 |
| *Oenanthe picata capistrata* | ZMUC, 29495 | Toepad | ThruPLEX DNA-Seq Kit | 96.7 | 5.3 |
| *Oenanthe picata opistholeuca* | ZMUC, 29578 | Toepad | ThruPLEX DNA-Seq Kit | 97.1 | 5.2 |
| *Oenanthe picata picata* | IR-TAN-005 | Blood | ThruPLEX DNA-Seq Kit | 98.8 | 14.2 |
| *Oenanthe pileata* | TCWC, 15606 | Muscle | Illumina DNA Prep Kit | 98.9 | 13.4 |
| *Oenanthe pleschanka* | CN-XS-006 | Blood | Chromium Genome Library kit | 99.8 | 15.1 |
| *Oenanthe seebohmi* | KA69373 | Blood | ThruPLEX DNA-Seq Kit | 99.2 | 11.2 |
| *Oenanthe warriae* | 12c | Blood | ThruPLEX DNA-Seq Kit | 98.7 | 13.4 |
| *Oenanthe xanthoprymna* | NHMO, 15188 | Blood | ThruPLEX DNA-Seq Kit | 99.0 | 12.9 |
| *Pinarochroa sordida* | YPM, ORN80066 | Toepad | ThruPLEX DNA-Seq Kit | 95.4 | 6.5 |
| *Thamnolaea cinnamomeiventris* | NRM, 20086147 | Muscle | ThruPLEX DNA-Seq Kit | 98.6 | 15.9 |

[a] AMNH; American Museum of Natural History; NHMUK: Natural History Museum, Tring; FMNH: Field Museum of Natural History; LACM: Natural History Museum of Los Angeles County; MVZ: Museum of Vertebrate Zoology, UC Berkeley; NHMO: Natural History Museum, University of Oslo; NRM: Naturhistoriska riksmuseet, Stockholm; TCWC: Texas A&M University Biodiversity Research and Teaching Collections; UWBM: University of Washington Burke Museum; YPM: Yale Peabody Museum; ZMUC: Zoological Museum, Natural History Museum of Denmark; ZFMK: Zoologisches Forschungsmuseum König. Samples for which no institution is indicated are part of the research group’s collection.

**Table S2 |** Number of loci across the genome using different filtering settings for 10 kb non-overlapping windows: minimum read depth (DP), minimum percentage of the window covered by data (PW), and missing data per site (MD).

| Filtering setting | Number of total windows | Number of no intra-locus recombination windows | Number of no intra-locus and free inter-locus recombination windows |
|---|---|---|---|
| DP=1, PW=50%, MD=15% | 29,730 | 10,051 | 6,791 |
| DP=5, PW=50%, MD=15% | 19,062 | 6,701 | 5,788 |
| DP=1, PW=50%, MD=5% | 26,030 | 9,600 | 6,476 |
| DP=1, PW=80%, MD=10% | 22,652 | 6,970 | 5,267 |

**Table S3 |** Selection of the best models among the maximum pseudo-likelihood networks in phyloNet 3.6.9.

| **Complex** | **Gene trees** | **Inferred admixtures** | **Likelihood** | **BIC** |
|---|---|---|---|---|
| *O. picata* | 7,310 | 0 | -33,633.58 | 67,278.88 |
| | | **1** | **-33,559.56** | **67,131.21** |
| | | 2 | -33,569.10 | 67,150.56 |
| | | 3 | -33,617.49 | 67,247.56 |
| | | 4 | -33,979.37 | 67,971.48 |
| | | 5 | -33,982.37 | 67,977.62 |
| *O. hispanica* | 7,335 | 0 | -33,326.92 | 66,665.57 |
| | | 1 | -33,180.89 | 66,373.86 |
| | | **2** | **-33,153.96** | **66,320.28** |
| | | 3 | -33,265.29 | 66,543.14 |
| | | 4 | -33,276.89 | 66,566.51 |
| | | 5 | -33,637.19 | 67,287.26 |
| *O. lugens* | ,7323 | 0 | -86,457.1 | 17,2926.3 |
| | | 1 | -85,643.3 | 17,1299.0 |
| | | **2** | **-85,014.9** | **17,0042.3** |
| | | 3 | -86,498.8 | 17,3010.4 |
| | | 4 | -86,574.7 | 17,3162.3 |
| | | 5 | -86,095.2 | 17,2203.3 |

### Supplementary Figures

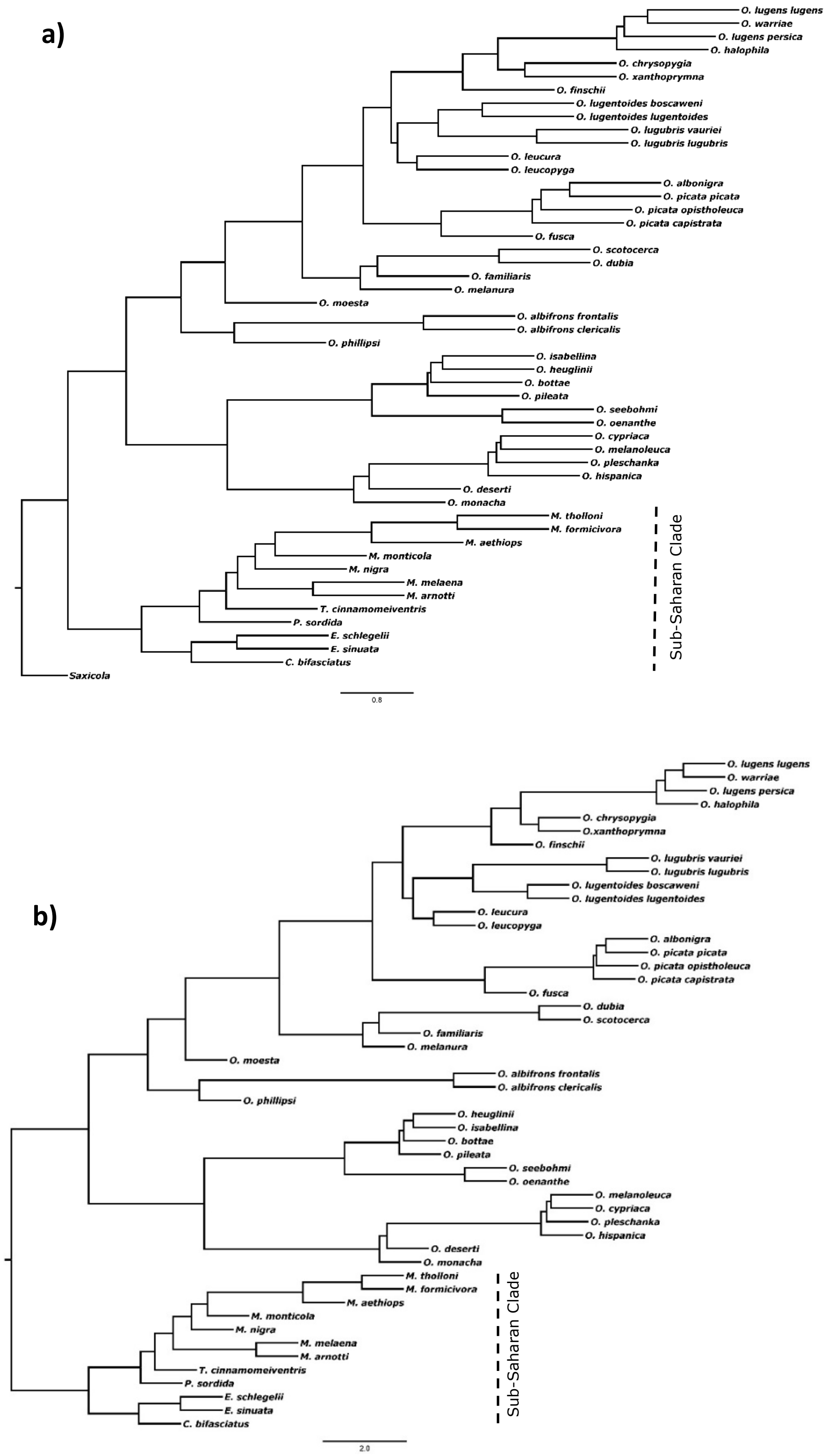

**Figure S1 |** Genome-wide multispecies coalescent species tree of open-habitat chats estimated in ASTRAL-III. All nodes have 1.0 local posterior probabilities. a) Species tree based on 2,091 BUSCOs, using *Saxicola* as an outgroup. b) Species tree based on 5,788 loci (non-overlapping 10 kb windows across the genome) using the sub-Saharan clade as an outgroup.

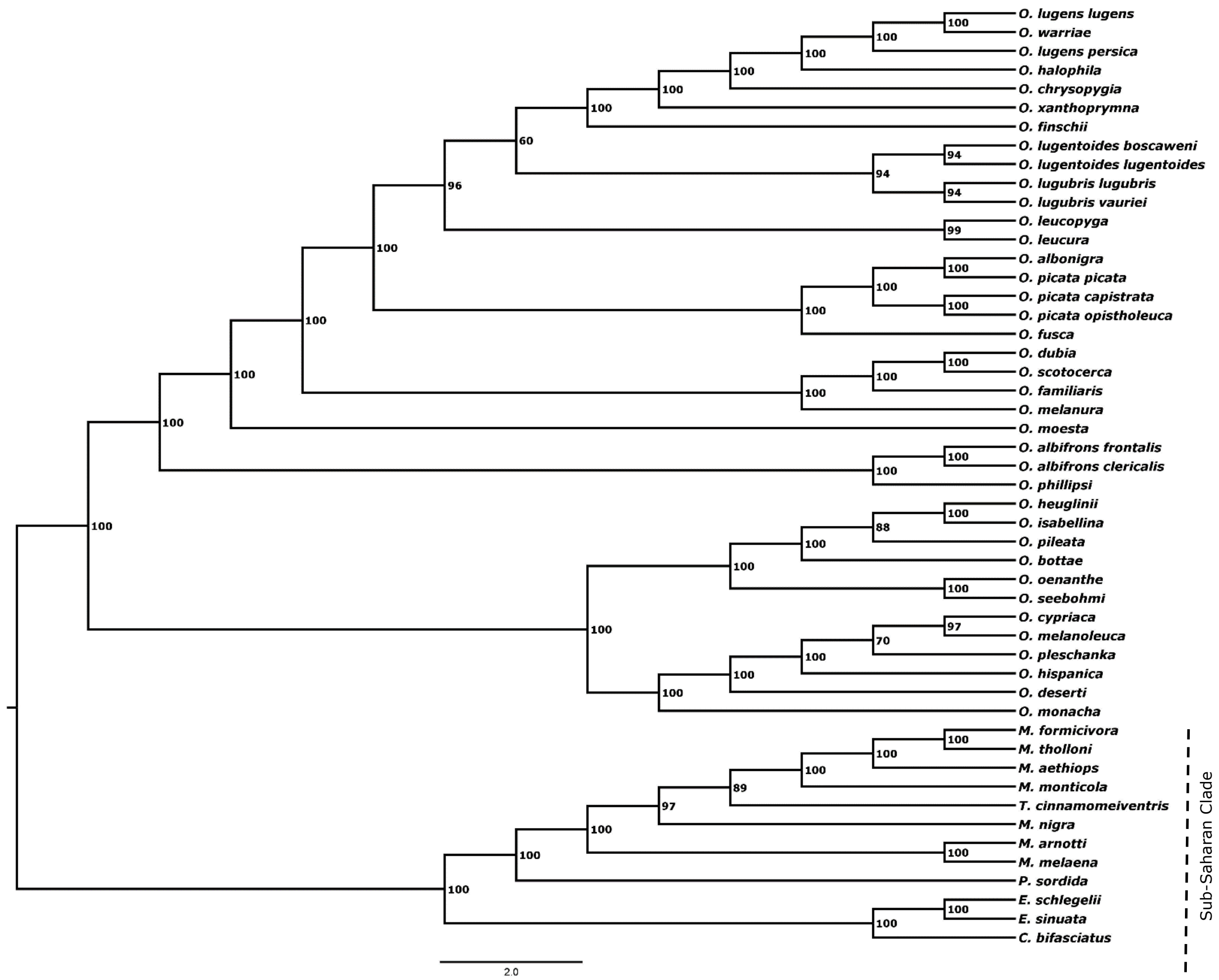

**Figure S2 |** SNP-based multispecies coalescent species tree of open-habitat chats inferred with SVDquartets in PAUP* 4. Maximum likelihood bootstrap support values are shown above branches.

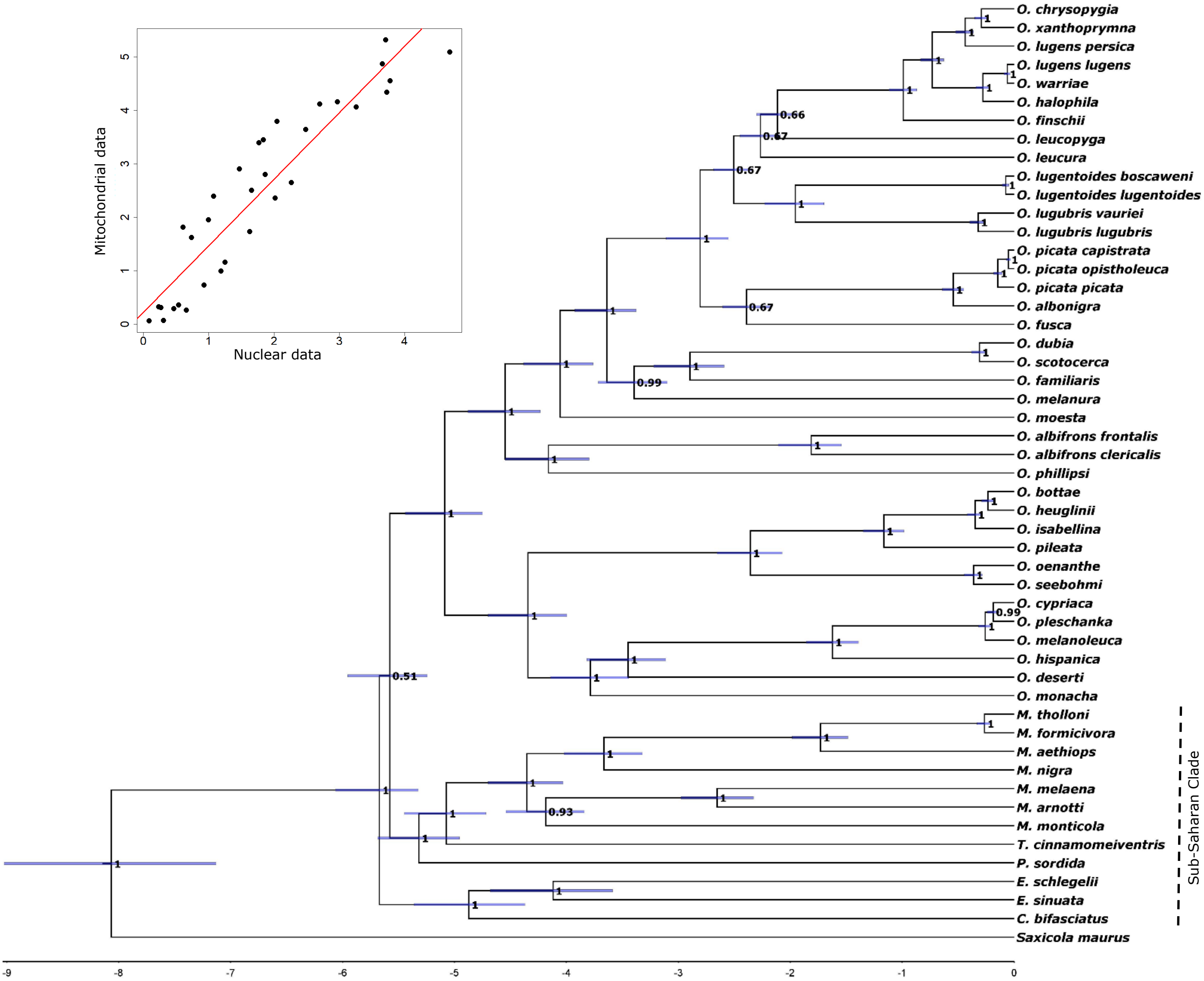

**Figure S3 |** Time-calibrated maximum clade credibility tree of the BEAST 2.6.6 analyses based on 13 protein-coding mitochondrial genes of open-habitat chats. Blue bars represent the 95% highest posterior density (HPD) distributions for the estimated divergence times. Posterior probabilities are indicated at nodes. The correlation between the dating based on the nuclear (BUSCOs) and mitochondrial data is shown at the top left (Pearson's r=0.93, p<0.001).

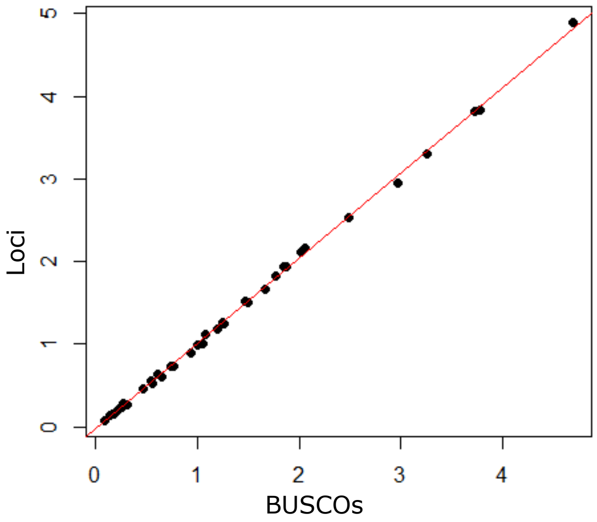

**Figure S4 |** Correlation between the time calibration based on 1.8 Mb high confidence BUSCOs and 3.8 Mb high confidence loci (10 kb non-overlapping windows across the genome) using RelTime-ML implemented in MEGA 11 (Pearson's r=0.99, p<2.2e-16).

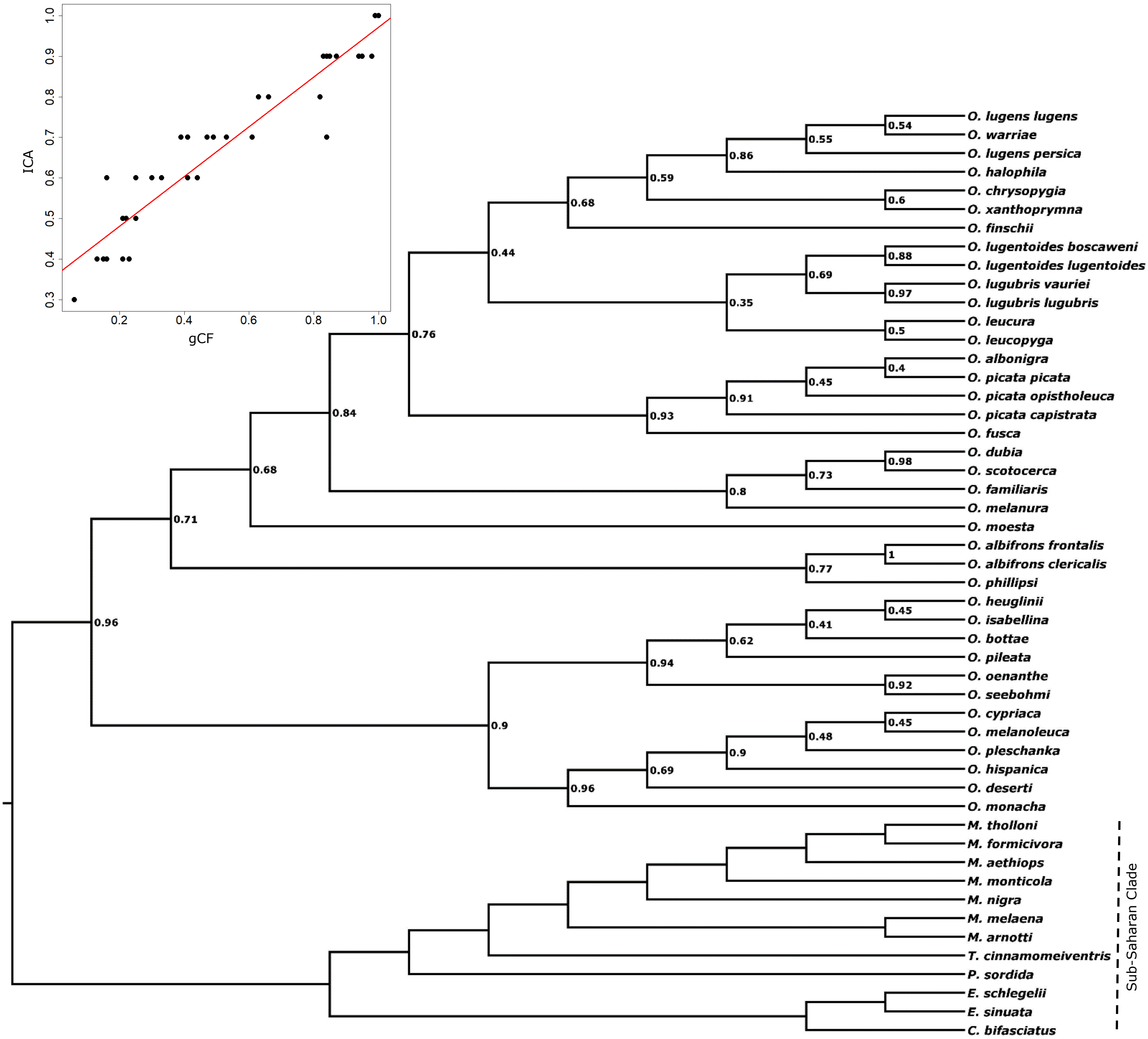

**Figure S5 |** Genome-wide multispecies coalescent species tree of open-habitat chats estimated in ASTRAL-III. ICA values for 29,730 maximum-likelihood gene trees are shown for each node. The high correlation between the gCF values calculated in IQ-TREE 2.1.2 and ICA values calculated in PhyParts 0.0.1 is shown at the top left (Pearson's r=0.94, p<0.001).
